# Representation of conceptual affordances in eye movements and the entorhinal cortex

**DOI:** 10.1101/2025.01.21.633955

**Authors:** Alexander Eperon, Christian F. Doeller, Stephanie Theves, Roberto Bottini

## Abstract

Structural representations in the entorhinal cortex are a crucial feature of cognitive maps. However, a complete understanding of task structure often requires knowledge of the actions available at each state (analogous to the moves of different pieces on a chessboard). Moreover, action representations are key to several models of the hippocampal-entorhinal system. In this study, we tested whether the entorhinal cortex represents the actions that allow transitioning between states of a conceptual space. In particular, participants learned to transition between numerically-labelled states using different mathematical operations. Behaviourally, participants learned and generalised action information across states. Action information was also reflected in eye movements, indicating the active processing of the possible actions for a given state during the trials. Importantly, we found that neural pattern similarity in the right entorhinal cortex scaled with the similarity of action affordances across the states. This affordance representation was not explained by other properties of the task space such as link distance between the states or action magnitude. In sum, this study provides first evidence for the integration of action information into ocular and entorhinal representations of conceptual spaces, suggesting that these may not just store experiences, but provide information about how to explore knowledge.

## INTRODUCTION

Human representations of space are closely connected to action. Whereas early maps tended to highlight topography useful to hunter-gatherers, modern mapping apps accentuate the differences between major roads and cycle paths.

The mammalian hippocampal formation is known to contain a variety of spatially-tuned cell types which respond selectively to locations in space (e.g. O’Keefe & Dostrovsky, 1971; Hafting et al., 2005; Høydal et al., 2019) and are typically thought to offer a neural substrate for a ‘cognitive map’ (Tolman, 1948; O’Keefe & Nadel, 1978). Indeed, recent evidence has supported the longstanding idea that the hippocampal formation encodes relations in support of memory, across spatial and non-spatial domains (Manns & Eichenbaum, 2006). For instance, studies have shown that navigation in a two-dimensional space spanned by non-spatial dimensions elicits a hexa-directional modulation of neural signals in the entorhinal cortex by the trajectory angle, mirroring the hexagonal grid pattern observed during navigation (Constantinescu et al., 2016; Viganò & Piazza, 2020; Park et al., 2021; Nitsch et al., 2023; Qasim et al., 2023; Schäfer et al., 2024). Moreover, the hippocampal formation also encodes other metrics of distance or position in abstract spaces, such as link distance on a graph (Schapiro et al., 2012; Garvert et al., 2017, 2023; Pudhiyidath et al., 2021), distances in conceptual and social spaces (Theves et al., 2019, 2020; Park et al., 2021; Schafer et al., 2022), or location along a sound frequency spectrum (Aronov et al., 2017).

To exploit relational knowledge, however, we need to understand how to transition between states - in reinforcement learning terminology, to take ’actions’ (Sutton & Barto, 1998). For instance, to win a chess game, we need to know which actions can move the pieces from the initial setup to checkmate. Indeed, action input is integral to several influential models of the hippocampal formation as the cue to move between different states (which might be memories or positions), whether built around a learned graph (Stachenfeld et al., 2017; Whittington et al., 2020; George et al., 2021) or a pre-structured Euclidean space (Chandra et al., 2025; for reviews of models see Dong & Fiete, 2024; Whittington et al., 2022). However, whether and how action information is actually incorporated in relational memory representations, i.e. in the entorhinal cortex, remains to be established empirically (Burak & Fiete, 2006; Ginosar et al., 2023; Morris & Derdikman, 2023).

In the context of spatial navigation, although the exact nature of action representation is unclear (Ginosar et al., 2023), there is increasing evidence that entorhinal representations are sensitive to changes in idiothetic signals (such as speed (Kropff et al., 2015, 2021) and gaze and head position (Mallory et al., 2021)), as well as behaviourally-relevant environmental changes. Specifically, entorhinal cells are sensitive to changes in landmarks or goal position (Boccara et al., 2019; Butler et al., 2019; Gonzalez & Giocomo, 2022; Wen et al., 2024). This has also been observed in human neuroimaging studies, which have shown distortions of grid-like signatures in fMRI based on goal locations (Viganò et al., 2023), arena layout (He & Brown, 2019) or prior levels of visual experience (Sigismondi et al., 2024). These distortions might be interpreted as a representation of ‘action-relevant space’ (Ginosar et al., 2023). Moreover, incorporating a condition of ‘actionability’ (that is, representations must enable actions to be carried out) appears to improve on existing models of grid cells by replicating key features such as multiple modules (Dorrell et al., 2022), and it has also been suggested that object or goal vector cells in the medial entorhinal cortex offer a neural substrate for action (Whittington et al., 2022). These studies point in the direction of entorhinal representations which are shaped and responsive to behaviour, but it remains an open question if action itself is represented in the entorhinal cortex independently of specific stimulus features or environmental layout.

Based on this, we ask whether the entorhinal cortex represents the possible actions from a given state. As with prior reports, we note the similarity between this idea and Gibson’s theory of affordances (Gibson, 1979; Behrens et al., 2018; Summerfield, 2022). To distinguish from motor affordances, however, we refer to the idea that the brain encodes the possible transitions (i.e. actions) in an abstract state space representing a graph of conceptual knowledge here as conceptual affordances.

In this study, we investigate the neural representation of conceptual affordances using fMRI and eye-tracking while participants evaluated numerical operations (actions) afforded by given numbers in a repeating graph structure (states). We found affordances to be represented in eye movements and entorhinal cortex activation patterns.

## RESULTS

### Learning and generalisation of state-action associations

We predicted that if the entorhinal cortex represents transition information in a conceptual space, neural pattern similarities would scale with the action possibilities afforded by different states. Firstly, therefore, we aimed to teach participants a repeating graph consisting of states and actions along a numberline: participants learned that certain numbers permitted transitions to other numbers (according to a predetermined graph structure, see Figure 1b) by free exploration along the graph using button presses. For instance, state 1, which could be instantiated by different numbers, afforded +2 or -2, while state 2 afforded +1 or +2. Knowledge of the graph was tested using blocked test trials presenting two candidate successors, of which only one corresponded to a possible transition and had to be chosen. In two subsequent sessions using the eyetracker and in the MRI scanner, respectively, participants were presented with individual numbers in a pseudorandom order (Figure 2b top). In these sessions their knowledge of the graph was probed using probe trials showing two numbers, to which participants had to respond if both were the correct successors for the previously-presented number or not (Figure 2b bottom) .

**Figure 1:**
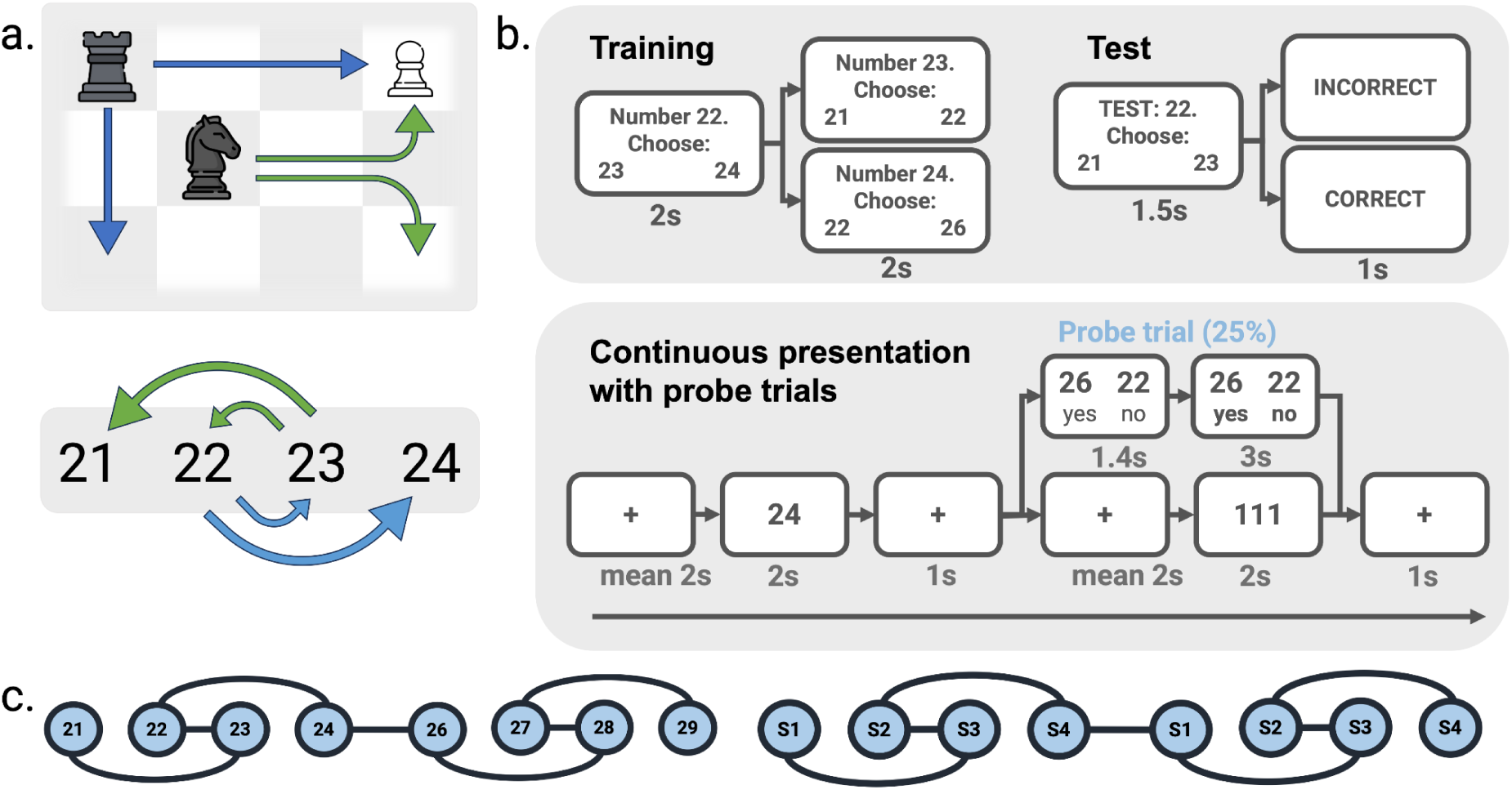
Experimental design. a. Representations of (conceptual) space may benefit from the integration of action affordances, e.g in a game of chess, where information about how pieces can move is necessary to play the game. We tested this idea using numerical operations. b. Participants learned how numbers may ’transform’ into other numbers using an exploration task, whereby they were presented with an individual number and were offered a free choice of two possible successors. When a successor was chosen, the next trial offered the successors for the chosen number. In interleaved test blocks, participants indicated which were the ’correct’ successors for each number. In the eyetracker and fMRI scanner, individual numbers were presented in a pseudorandom order. To ensure attention on the learned actions, one quarter (16) of trials were followed by probe trials in which participants indicated if two presented numbers were the correct successors. c. Unbeknownst to participants, number-action bindings formed part of a repeating sequence arranged in graph ’modules’ of four numbers. This can be seen as a set of four states with the same action affordances. As the modules repeated every five numbers, the structure allowed easy inference of action possibilities for any new numbers. Chess images adapted from flaticon.com.

**Figure 2:**
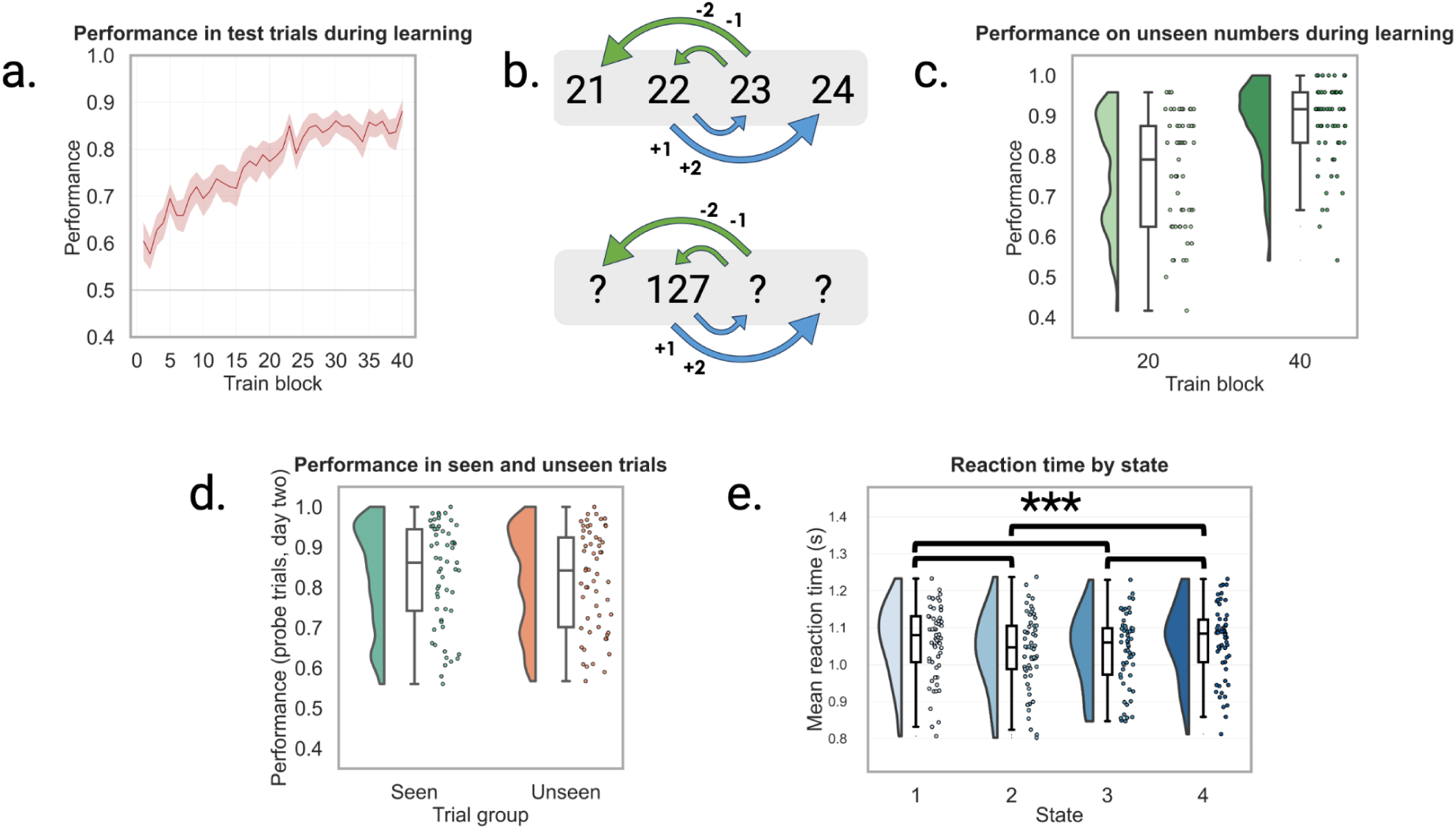
Behavioural performance. a. Participants learned the numbers and successors well, achieving a performance of 84.7% by the end of the first day). b. The repeating structure should allow participants to infer the correct successors for previously-unseen numbers. c. Halfway through the first day’s training, and at the end of each training session, participants were presented with a longer test block with new numbers and no feedback. After 20 blocks performance was at mean 76.2%, but by the end of training they reached a performance level of 88.8%. d. In the recall task in the fMRI scanner, participants were able to correctly identify the successors when presented with unexpected probe trials (mean performance 82.7%). There was no difference in performance between previously-seen numbers and new numbers in the scanner task (p=0.126). e. Reaction times showed faster responses for states 2 and 3 relative to states 1 and 4. Asterisks indicate significance.

Participants were trained on two consecutive days to learn how specific numbers could ’transform’ into other numbers. By the end of the first day, participants were able to reliably select the correct transitions for different numbers (accuracy: 84.7%; chance level 50%, std. 7.1%, last ten blocks; figure 1a). Moreover, they were able to apply the learned transformations to new numbers: after training on the second day, they reached a performance level of 88.8% on unseen numbers (std. 11.3%). The application of these operations to hitherto-unseen numbers indicates that participants understood that numbers and transitions were factorised.

In the recall task on the first day (during eyetracking), performance was lower than in the previous task (accuracy: mean 68.1%, std. 15.9% across day 1; missed responses: mean 8.4%, std. 6.4%), but nonetheless improved as familiarity with the task increased to 71.5% in the fourth and final block (std. 21.6%; difference between fourth and first block T=218.5, p=0.006, Wilcoxon signed rank test). Participants were better at answering correctly for seen than unseen numbers (seen: mean 70.8%; unseen: mean 65.8%; t(42)=2.78; p=0.008; see Figure 2c).

Knowledge of the structure persisted to the second day, suggesting a long-term encoding of the graph information. In the MRI scanner, performance was at a mean accuracy of 82.7% (std. 12.5%; missed responses: mean 7.8%, std. 6.3%). Moreover, the differences between seen and unseen numbers diminished to a non-significant level, suggesting that longer-term learning was dominated by the abstract structure rather than number-specific action bindings (no significant difference between seen and unseen numbers: seen: mean 83.2%, std. 12.8%; unseen: mean 82.2%, std. 12.3%; t(56) = 1.56, p=0.13; see Figure 2d).

A stimulus-independent encoding of the graph structure was supported by state-specific reaction time differences during probe trials. Reaction time data in the recall task on the second day showed a difference between response times for different states: state identity predicts reaction times (F(3, 168)=9.77, p<0.0001; see Figure 2e). Post-hoc pairwise comparisons show a significant difference between states 1 and 2 (t(56)=4.44, p<0.0001), states 1 and 3 (t(56=3.45, p=0.0011), states 2 and 4 (t(56)=-4.09, p=0.0001) and states 3 and 4 (t(56)=-3.38, p=0.0014). There were no significant differences between states 1 and 4 (t(56)=0.17, p=89) and states 2 and 3 (df(56)=-0.23, p=0.82).

Nonetheless, the majority of participants did not self-report reliance on the learned action affordances, despite their necessity to solve the task for unseen numbers. Questionnaire data indicated a mix of strategies (specifically, 42.1% reported memory-based strategies, 24.5% relied on calculation and the remaining 33.3% used both), although no participants were aware of the experimental question. This indicates that although participants needed to learn actions in order to solve the task, they were not necessarily aware of a process of abstraction and then calculation.

In short, participants were able to learn the numerical operations afforded by different numbers, and apply this knowledge to new numbers. This indicated knowledge of a repeating graph consisting of state-action bindings.

### Eye movement direction reflects the sign of conceptual affordances

Prior work has suggested that exploration of conceptual spaces is reflected in eye movements (Viganò et al., 2024). As previous evidence has shown a clear lateralisation of responses to numerical stimuli, predicted that if participants engage with the task using a mental numberline, eye movements would skew left or right based on the type of actions allowed. In particular, we predicted that eye movements would skew more rightwards for states that allow only positive actions (e.g. state 2) relative to states that allow only negative actions (e.g. state 3).

We compared the relative gaze position difference for these states for each timepoint in a [-0.5,2.5] window around stimulus onset and tested for each timepoint in the trial if movement shift was greater than 0. There was a significant effect in both the x-axis (t(42)=5.99, p<0.001, cluster window of 845 to 1778 ms after onset and t(42)=3.83, p=0.002, cluster window of 1873 to 2200 ms after onset, 14 clusters; Figure 3b), as well as the y-axis (t(42)=3.17, p=0.018, cluster window of 1139 to 1423 ms after onset, 16 clusters; Figure 3c). Finally, we show that ocular responses were skewed more strongly left-or-rightwards in better-performing participants on the x-axis (r=0.488, p=0.00090, n=43; Figure 3d), but not on the y-axis (r=0.0830, p=0.597 in y axis; n=43). This indicates that a stronger (horizontal) schematic implementation (i.e. mental numberline) may be useful to carrying out the task.

**Figure 3:**
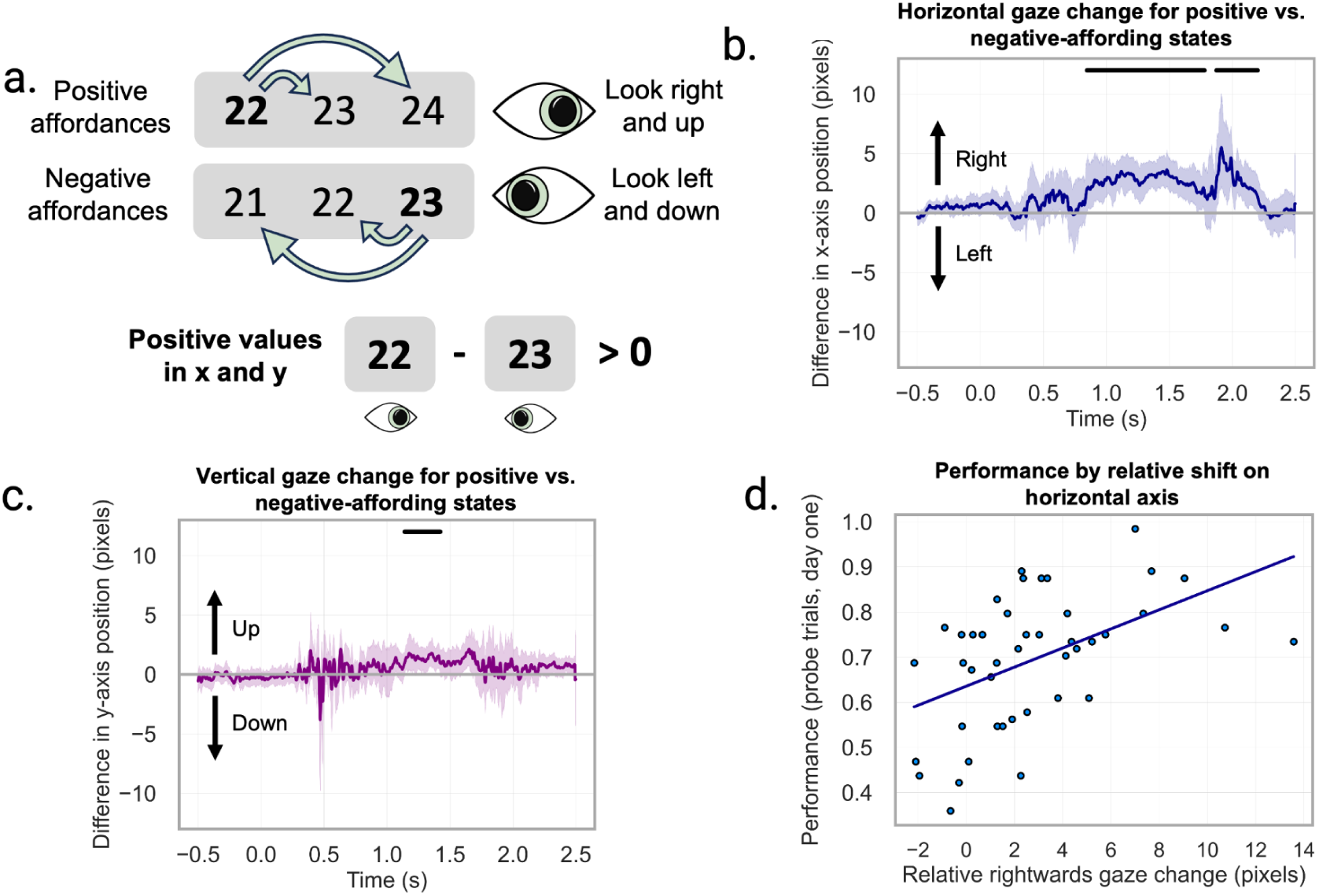
Ocular representation of conceptual affordances. a. We predicted that, if participants were employing a mental numberline, eye movements would skew further right and upward for numbers which afford positive actions, relative to numbers which afford negative actions. b., c. Relative fixations for states 2 and 3. When the mean position of state 3 (which is mapped to number 23 in the example) was subtracted from the mean position for state 2 (mapped to number 22), we find a positive value in both the x and y axis, indicating a more positive shift for more positive actions. Horizontal bars indicate significance. d. The extent of the effect in the x-axis (median position change during the cluster window) correlated with performance in the probe trials of the recall task during eye-tracking (r=0.49, p<0.001).

In sum, while the behavioural results of the learning and test phase (Figure 1b top) shows that participants had learned and generalised the affordances across different instances of the states, their reflection in eye movements further indicates that they actively processed affordance information when cued with a given state during the measurement trials (Figure 1b bottom). Thus, next we approached our main question of whether the entorhinal cortex would represent states in terms of the actions they afford.

### Entorhinal pattern similarity reflects affordances of states

The main prediction was that the entorhinal cortex would represent conceptual affordances; that is, neural responses would be more similar when states allowed similar actions. To test this, we used representational similarity analysis to compare neural pattern similarity across the states to a model RDM indicating the number of shared possible actions between states. Indeed, we found a representation of conceptual affordances in the right entorhinal cortex (right: t(56)=2.13, p=0.019; left: t(56)=-0.49, p=0.69). We further assured that this result is not driven by other effects related to affordance magnitude (the unsigned magnitude of afforded actions) or link distance, by including these alternative model RDMs (see Methods; see model RDMs in Figure 4c) as covariates and replicating the finding (right: t(56)=2.29, p=0.013; left: t(56)=-0.60, p=0.73; Figure 4a-c).

**Figure 4:**
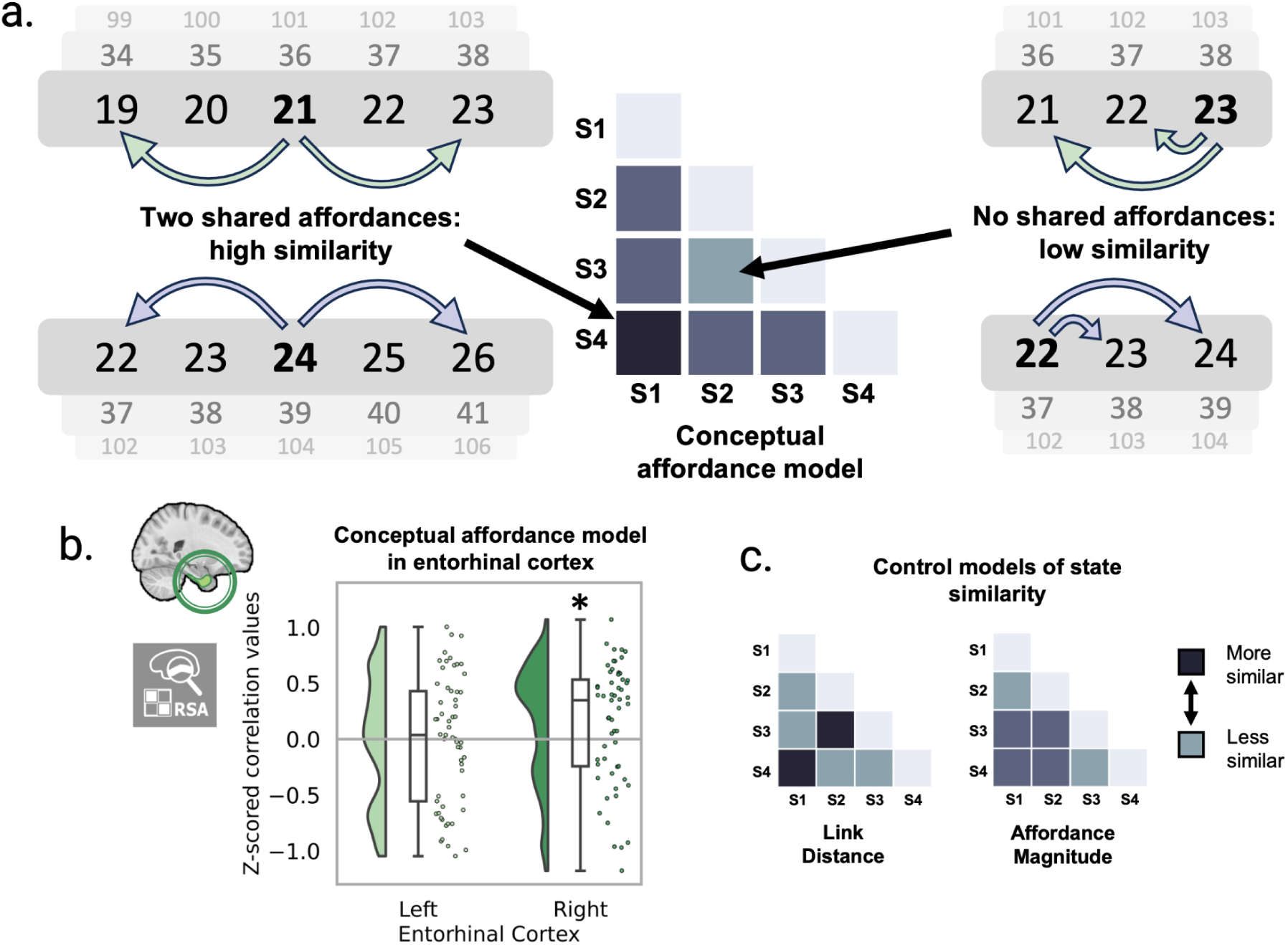
Neural representation of conceptual affordances. a. We used representational similarity analysis to compare neural pattern similarity across states in the entorhinal cortex with a model of state similarity based on the number of shared affordances. b. Neural pattern dissimilarity matrices (neural RDMs) were calculated in an a prior region-of-interest (ROI) using cross-validated Mahalanobis distance. Neural patterns in the right entorhinal cortex were correlated with the conceptual affordances RDM. c. Control models of state similarity. Asterisks indicate significance.

To further confirm the stability of the entorhinal affordance representation, we confirmed that cross-validated mean distances in the entorhinal RDMs were greater than 0, indicating our neural RDMs reflected consistent differences in neural responses to different conditions across runs. The affordance effect persisted within a subset of subjects whose mean RDM values were greater than 0, indicating that for subjects in which entorhinal voxels respond consistently to different states, neural patterns correlate with the predicted affordance model. (Supplementary Figure 1). Moreover, we used a novel variant of reliability-based voxel selection (Tarhan & Konkle, 2020) to sub-sample voxels within our ROIs which responded consistently across different runs. Likewise, we replicated the affordance representation in the entorhinal cortex, indicating that it is driven by reliable, i.e. more condition-responsive, voxels (Supplementary Figure 2). Finally, the effect is not below the expected noise ceiling in the right entorhinal cortex, and therefore as strong as it could be given noise (Supplementary Figure 3). For details, see Supplementary Materials.

In sum, we find that the entorhinal cortex represents the conceptual affordances per state. This effect is not driven by alternative properties of the structure such as link distance or the affordance magnitude and is stable with respect to selected participants, voxels and noise ceiling.

## DISCUSSION

The hippocampal formation has a demonstrated role in mapping out relational knowledge (O’Keefe & Nadel, 1978; Behrens et al., 2018; Bellmund et al., 2018; Bottini & Doeller, 2020) with potential implications for general cognitive performance (Tenderra & Theves, 2024). In the absence of idiothetic signals, however, it is unclear how the brain ’navigates’ these memory representations - to play a game of chess we need to know both the layout of the pieces and how they move. Prior work has demonstrated that the entorhinal cortex represents action-relevant information during spatial navigation. Here we provide explicit evidence in humans that the entorhinal cortex represents actions in a conceptual space.

We used a numberline as base structure to which states with different conceptual affordances were associated, thereby allowing us to distinguish action representations from other features of a state space (e.g. state identity, link distance). We find that the entorhinal cortex represents the actions ’afforded’ by a given state.

In line with previous work linking eye movements to a simulated mental numberline (Dehaene et al., 1993; Hartmann, 2022; Viganò et al., 2023, 2024), conceptual affordances were also tracked by eye movements: participants looked further rightwards when afforded actions were positive relative to states with negative action affordances. Compared to previous work, we show that the gaze effect is positively correlated with task performance, suggesting a functional role for attention-based navigation in solving the task (Giari et al., 2023).

Models of hippocampal processing that are based around reinforcement learning such as the Tolman-Eichenbaum Machine (TEM) (Whittington et al., 2020) or Successor Representation (Stachenfeld et al., 2017), often consider action input to move between states. In the context of the hippocampal formation, this could potentially be seen as a non-spatial abstraction of the velocity input required by models built around the navigation of Euclidean spaces (e.g. vector-hash, (Chandra et al., 2025)). While in physical navigation velocity input may be calculated from idiothetic information, in non-spatial domains action poses more of a conundrum. The present study indicates that information regarding action identity is present in the entorhinal cortex, although establishing the exact nature of the signal will require further study.

More generally, our work fits in with a growing body of evidence that cognitive maps are closely tied to learned behaviour. Recent work in humans has shown that neural representations in the entorhinal cortex and hippocampus are influenced by task demands or the relevance of dimensions and goals (Theves et al., 2020; Crivelli-Decker et al., 2023; Sweigart et al., 2023). Prior work has shown that when learning new concepts the hippocampus maps stimuli according to the conceptually-defining, but not all perceptual dimensions (Theves et al., 2020) and hippocampal and grid-like entorhinal representations of the conceptual space only emerge after relational encoding becomes task-relevant (Theves et al., 2020; Schäfer et al., 2024). Blind humans exhibit a four-directional symmetry instead of the typical hexadirectional symmetry likely elicited by grid cell populations, suggesting that entorhinal representations may reflect long-term behavioural patterns (Sigismondi et al., 2023). Similarly, rodent and human work has shown a distortion in grid signal as a function of goal placement (Boccara et al., 2019; Butler et al., 2019; Viganò et al., 2023). Moreover, rodent intracranial work has shown that grid cells display distortions around landmarks that appear to reflect future behaviour. These distortions are too slow to explain one-shot learning (the authors invoke a mediating plasticity mechanism as a potential solution), but nonetheless suggest a coupling of entorhinal representations with behaviour (Wen et al., 2024; for landmark alignment in a mental simulation see Hwang et al., 2023; Neupane et al., 2024). Our findings generally support the intuition that entorhinal representations are influenced by possible actions (for a computational model of grid cells based on action constraints, see Dorrell et al., 2022).

There are multiple possible neural substrates for processing of actions in the entorhinal-hippocampal system. In spatial navigation, action is integrated into the grid cell network via velocity input, which may be derived from idiothetic signals or internally generated (Burak & Fiete, 2006; Khona & Fiete, 2022; Dong & Fiete, 2024) and can be described by specialised cell types in the entorhinal cortex (Kropff et al., 2015; Mallory et al., 2021; Munn et al., 2020). Intriguingly, Mallory and colleagues (2021) identify entorhinal cells which conjunctively code for rodent eye and body position, reminiscent of saccade direction cells in head-fixed primates (Killian et al., 2015; Killian & Buffalo, 2018). Beyond spatial navigation, it has been suggested (Whittington et al., 2020, 2022) that action computations can be carried out through vector cells such as object-vector cells (Høydal et al., 2019) and goal-vector cells (Sarel et al., 2017). In light of the role for vision in primates and work potentially suggesting a role of eye movements in ‘simulating’ conceptual navigation (eye movements related to a grid-like signal: Killian & Buffalo, 2018; Nau et al., 2018; Staudigl et al., 2018; Giari et al., 2023; eye movements reflecting conceptual navigation: Viganò et al., 2023, 2024), our findings could imply that saccade cells may play a role similar to that proposed for vector cells in integrating action information during spatial navigation. Saccade direction cells in the primate entorhinal cortex encode the direction of eye movements during free viewing of complex images (Killian et al., 2015). Killian and colleagues compared these cells to head direction cells in rodents, because of their directional selectivity. However, it remains to be tested if these cells hold a vectorial representation (which may be calculated via the grid system (Bush et al., 2015; Bicanski & Burgess, 2019), encoding not only the direction but also the magnitude of movement.

The present results provide evidence that the entorhinal cortex represents conceptual affordances. These findings should lend weight to a close link between cognitive maps and behaviour. In his seminal work on visual perception, James Gibson eschewed the idea of cognitive maps because of the lack of an “internal perceiver to read [the] map”. (Gibson, 1979) Our work suggests that, on the contrary, cognitive maps can provide a normative guide to navigation through action representation.

## MATERIALS AND METHODS

### PARTICIPANTS

Sixty healthy German-speaking volunteers participated in the study. All participants were right-handed, with normal or normal-to-corrected vision and reported no history of neurological or psychiatric disorders. All participants provided written informed consent to participate in the study, and were compensated for participation. The study was approved by the local ethics committee at the University of Leipzig, Germany (protocol number 437/22-ek). Three participants did not reach a predetermined performance threshold of 74/128 correct answers in the scanner task, which corresponded to a performance greater than chance in a binomial test at an alpha level of 0.05. These participants were excluded from all analyses. Within the remaining fifty-seven subjects, 29 self-identified as female with an overall mean age of 26.7 (s.d. 4.9). In addition, a further 14 participants had to be excluded from the eye-tracking analysis due to difficulties with calibration or poor data quality identified during testing, resulting in a final sample of forty-three for ocular analyses (19 self-identified female; mean age 25.8, s.d. 4.8).

The sample size was determined without a prior power analysis, due to the difficulties in making an accurate estimate deriving from the small number of previous related studies and the multiple planned analyses. Initially, a high drop-out rate was expected due to a taxing task involving numbers, and therefore it was decided to recruit sixty participants in order to match or exceed the larger sample sizes typical of studies which have been able to detect comparable effects in the entorhinal cortex (Garvert et al., 2023; Nitsch et al., 2024; Viganò et al., 2023).

### EXPERIMENTAL DESIGN

The experiment took place over two consecutive days. On the first day, participants were informed that they would play the role of an archeologist who had discovered a new number system in local sandstone caves whereby numbers could be transformed into other numbers. This was, of course, fictitious and designed to ensure they understood that they should try to learn number transitions from scratch rather than apply any pre-existing intuitions. Participants were instructed that their task was to first understand how numbers could ’transform’ into other numbers, and then use this knowledge to identify correct number transformations.

Each day consisted of both training and a recall task. The training was designed to teach participants how numbers were connected to other numbers by interleaved exploration and test blocks (see below). The recall task was used to measure neural and ocular responses to each number.

On the first day, participants carried out the training for approximately an hour and a half (40 blocks), followed by half an hour of a recall task (4 runs) while eye movements were monitored using an eye tracker (details below). On the second day, they repeated the training for only 20 minutes (10 blocks), followed by an hour of the recall task (8 runs) while brain activity was monitored using fMRI. At the end of the experiment, participants were asked to fill out a questionnaire, which primarily aimed to test strategies used while carrying out the task, and gather relevant demographic information (see attached materials).

### METHOD DETAILS

#### Stimuli

Our design aimed to distinguish between previously-tested features of a state space (e.g. link distance) and the specific transitions (numerical operations, referred to interchangeably as ’actions’) that could be carried out within this structure.

To dissociate state identity from specific stimulus identity, we created a repeating structure (module) based on relations between four states. This module repeated every five numbers, allowing easy generalisation while avoiding visual confounds in the design. Each state (number) was associated with two possible actions (arithmetic calculations:+2,-2,+1,-1) and participants could navigate the numberline by choosing one of them. For instance, if number 3 afforded actions -1/-2, it would be possible to select 1 but not 5 as the next state.

Using a numberline as the base structure allowed us to manipulate the transitions independently from the specific states. For example, transitioning between any two numbers could involve a big numerical change but still constitute one link in the graph structure. As previous studies reported a linear encoding of numerical knowledge, we assumed that the relationship between actions would hold independently of the numbers seen (e.g. ’2’ to ’5’ would be similar to ’55’ to ’57’). Likewise, while a movement from ‘3’ to ‘2’ may be a single step, it constitutes a different transition (−1) from that of the next step ‘2’ to ‘4’ (transition of +2).

To further control for any pre-learned mathematical associations, we balanced number-state assignments across participants.

In total, participants were trained and tested on 4 and 8 modules (16 and 32 numbers) respectively prior to entering the fMRI scanner, while each run consisted of an extra 3 previously unseen modules (12 numbers). In total, the available structure seen throughout the entire experiment consisted of 36 modules per participant, corresponding to 144 numbers. All numbers were greater than 20, to avoid interactions with pre-existing associations related to single-digit numbers.

#### Training

Training was designed to teach participants the transformations between numbers using a ’free exploration’ paradigm. Participants were presented with one number on the screen, alongside two options which corresponded to the two directly connected numbers in the graph structure introduced above. Selecting a number from the two choices led to the next display including the selected number with its two direct connections (see fig. 1). This phase can be seen as an exploration of the numberline using the numerical operations available in the structure. After 16 exploration trials, participants were presented with 18 test trials on the same numbers, in which only one of the two options corresponded to a correct successor according to the graph structure.

After 20 and 40 of these train and test blocks, as well as at the end of the training on the second day, 24 test trials using novel numbers (drawn from the beginning and end of the participant-specific numberline) were presented without feedback. This allowed a test of how well participants applied the learned transitions to new numbers, which we refer to as ’weak generalisation’ due to remaining clues (e.g. last digit of the number stays the same).

Test trials were evenly selected from different positions within stimulus modules. Most trials were selected from within a range of two from the presented number in order to avoid use of a distance-based heuristic. Nonetheless, to ensure participants paid attention to the entire number rather than the last digit alone, each test block included two trials where the potential successors differed from a correct answer by a multiple of five.

#### Recall Task (eye tracker and MRI)

During eye-tracking and the fMRI session participants were presented with a series of numbers, including previously seen and new numbers. To keep participants engaged in the task, at random intervals probe trials queried their understanding of the affordance-state bindings.

If these two numbers were the two correct successors of the most recent number, participants had to press a button to ’accept’ them. Otherwise, they had to reject the option. Half of probe trials showed correct possible successors.

Buttons were counterbalanced across runs, and the ordering of presented probe trial stimuli was randomised, to ensure there was no systematic bias in either motor responses or visual attention. To avoid potential problems of temporal autocorrelation in the MRI scanner, the order of conditions was counterbalanced such that each trial was preceded an equal number of times by trials from different conditions.

Inter-stimulus (ISI) intervals were drawn from a truncated exponential distribution with a mean of 3s in the MRI scanner and 2.5s during eyetracking, respectively. To guarantee that the ISI selection did not influence the final results due to temporal autocorrelation, for each run we ensured that the Spearman correlation between ISI length between two trials and model-based trial distance was less than 0.1 via permutation of possible ISI sequences across all models (see ’Analysis: Representational Similarity Analysis’ for model details).

In the eyetracker, participants carried out four runs of approximately six minutes each. Each run consisted of 80 trials, of which 16 were probe trials.

In the MRI scanner, participants carried out eight runs, each of which lasted seven minutes. As with the eyetracker, each run consisted of 80 trials of which 16 were probe trials. This resulted in a total of 512 trials for analysis.

#### MRI Data Acquisition

MRI data were recorded using a 3 Tesla Siemens Magnetom Prisma Fit scanner (Siemens, Erlangen, Germany) with a 32-channel head coil.

Following a localiser scan, blood-oxygen-level-dependent (BOLD) contrast was measured for the eight runs of the recall task using T2*-weighted whole-brain gradient-echo planar imaging (GE-EPI) with a multiband acceleration factor of 5.

The fMRI sequence has the following parameters: TR = 1000ms; TE = 22ms; voxel size = 2.5mm isotropic; field of view = 204mm; flip angle = 62°; partial fourier = 0.75; bandwidth = 1794 Hz/Px; multi-band acceleration factor = 5; 65 slices interleaved; distance factor = 10%; phase-encoding direction: A -> P. Per run, 415 volumes were acquired.

After every run, fieldmaps were recorded as a way to measure and correct for inhomogeneities in the magnetic field. Fieldmaps were acquired using opposite phase-encoded EPIs, using the same voxel sizes and field of view with TR = 8000ms; TE = 50ms; flip angle = 90°; partial fourier = 0.75; bandwidth = 17.934 Hz/Px.

Following recommendations from previous work (Nau, 2022), the acquisition box was manually aligned by the experimenter to line up with the long axis of the hippocampus. This has been suggested as a method to acquire clearer signal in the medial temporal lobes.

After each scanning session, an anatomical scan was recorded. This was done using a T1-weighted MP2RAGE (TR = 5000ms, TE = 19.6ms, voxel size = 1 mm isotropic). The resulting scan was denoised using the method outlined in (O’Brien et al., 2013).

All stimuli were projected onto a screen situated above the participant using a mirror attached to the head coil. Behavioural responses were collected using an MRI-compatible button box.

#### Eyetracker Data Acquisition

Participants were seated in front of a monitor leaning on a chin rest and forehead bar, at a distance of 57cm from the screen. We recorded binocular gaze position continuously using an EyeLink 1000 Plus (SR Research), at a sampling rate of 1,000 Hz. To ensure accurate recording, we calibrated the system at the start of each run using the built-in calibration and validation protocols from the EyeLink software using nine fixation points.

#### Preprocessing of fMRI data

Preprocessing was carried out using fMRIprep 22.0.1 (see below for a boilerplate provided by fmriprep). The processing steps include segmentation, head motion estimation and correction, slice timing correction, co-registration to anatomical images, fieldmap-based (AP-PA) susceptibility distortion correction and resampling to MNI space. All analyses were carried out in volumetric space, using subject space (T1w) for all analyses carried out at the single-subject level. Before running general linear models (GLMs) for any analysis, volumetric data was smoothed using a Gaussian FWHM filter with a 5mm width.

Results included in this manuscript come from preprocessing performed using fMRIPrep 22.0.1 (Esteban, Markiewicz, et al. (2018); Esteban, Blair, et al. (2018); RRID:SCR_016216), which is based on Nipype 1.8.4 (K. Gorgolewski et al. (2011); K. J. Gorgolewski et al. (2018); RRID:SCR_002502).

#### Preprocessing of B0 inhomogeneity mappings

A total of 8 fieldmaps were found available within the input BIDS structure for each subject. A B0-nonuniformity map (or fieldmap) was estimated based on two (or more) echo-planar imaging (EPI) references with topup (Andersson, Skare, and Ashburner (2003); FSL 6.0.5.1:57b01774).

#### Anatomical data preprocessing

A total of 1 T1-weighted (T1w) images were found within the input BIDS dataset.The T1-weighted (T1w) image was corrected for intensity non-uniformity (INU) with N4BiasFieldCorrection (Tustison et al. 2010), distributed with ANTs 2.3.3 (Avants et al. 2008, RRID:SCR_004757), and used as T1w-reference throughout the workflow. The T1w-reference was then skull-stripped with a Nipype implementation of the antsBrainExtraction.sh workflow (from ANTs), using OASIS30ANTs as target template. Brain tissue segmentation of cerebrospinal fluid (CSF), white-matter (WM) and gray-matter (GM) was performed on the brain-extracted T1w using fast (FSL 6.0.5.1:57b01774, RRID:SCR_002823, Zhang, Brady, and Smith 2001). Volume-based spatial normalization to two standard spaces (MNI152NLin6Asym, MNI152NLin2009cAsym) was performed through nonlinear registration with antsRegistration (ANTs 2.3.3), using brain-extracted versions of both T1w reference and the T1w template. The following templates were selected for spatial normalization: FSL’s MNI ICBM 152 non-linear 6th Generation Asymmetric Average Brain Stereotaxic Registration Model [Evans et al. (2012), RRID:SCR_002823; TemplateFlow ID: MNI152NLin6Asym], ICBM 152 Nonlinear Asymmetrical template version 2009c [Fonov et al. (2009), RRID:SCR_008796; TemplateFlow ID: MNI152NLin2009cAsym].

#### Functional data preprocessing

For each of the 8 BOLD runs found per subject (across all tasks and sessions), the following preprocessing was performed. First, a reference volume and its skull-stripped version were generated using a custom methodology of fMRIPrep. Head-motion parameters with respect to the BOLD reference (transformation matrices, and six corresponding rotation and translation parameters) are estimated before any spatiotemporal filtering using mcflirt (FSL 6.0.5.1:57b01774, Jenkinson et al. 2002). The estimated fieldmap was then aligned with rigid-registration to the target EPI (echo-planar imaging) reference run. The field coefficients were mapped on to the reference EPI using the transform. BOLD runs were slice-time corrected to 0.452s (0.5 of slice acquisition range 0s-0.905s) using 3dTshift from AFNI (Cox and Hyde 1997, RRID:SCR_005927). The BOLD reference was then co-registered to the T1w reference using mri_coreg (FreeSurfer) followed by flirt (FSL 6.0.5.1:57b01774, Jenkinson and Smith 2001) with the boundary-based registration (Greve and Fischl 2009) cost-function. Co-registration was configured with six degrees of freedom. Several confounding time-series were calculated based on the preprocessed BOLD: framewise displacement (FD), DVARS and three region-wise global signals. FD was computed using two formulations following Power (absolute sum of relative motions, Power et al. (2014)) and Jenkinson (relative root mean square displacement between affines, Jenkinson et al. (2002)). FD and DVARS are calculated for each functional run, both using their implementations in Nipype (following the definitions by Power et al. 2014). The three global signals are extracted within the CSF, the WM, and the whole-brain masks. Additionally, a set of physiological regressors were extracted to allow for component-based noise correction (CompCor, Behzadi et al. 2007). Principal components are estimated after high-pass filtering the preprocessed BOLD time-series (using a discrete cosine filter with 128s cut-off) for the two CompCor variants: temporal (tCompCor) and anatomical (aCompCor). tCompCor components are then calculated from the top 2% variable voxels within the brain mask. For aCompCor, three probabilistic masks (CSF, WM and combined CSF+WM) are generated in anatomical space. The implementation differs from that of Behzadi et al. in that instead of eroding the masks by 2 pixels on BOLD space, a mask of pixels that likely contain a volume fraction of GM is subtracted from the aCompCor masks. This mask is obtained by thresholding the corresponding partial volume map at 0.05, and it ensures components are not extracted from voxels containing a minimal fraction of GM. Finally, these masks are resampled into BOLD space and binarized by thresholding at 0.99 (as in the original implementation). Components are also calculated separately within the WM and CSF masks. For each CompCor decomposition, the k components with the largest singular values are retained, such that the retained components’ time series are sufficient to explain 50 percent of variance across the nuisance mask (CSF, WM, combined, or temporal). The remaining components are dropped from consideration. The head-motion estimates calculated in the correction step were also placed within the corresponding confounds file. The confound time series derived from head motion estimates and global signals were expanded with the inclusion of temporal derivatives and quadratic terms for each (Satterthwaite et al. 2013). Frames that exceeded a threshold of 0.5 mm FD or 1.5 standardized DVARS were annotated as motion outliers. Additional nuisance timeseries are calculated by means of principal components analysis of the signal found within a thin band (crown) of voxels around the edge of the brain, as proposed by (Patriat, Reynolds, and Birn 2017). The BOLD time-series were resampled into standard space, generating a preprocessed BOLD run in MNI152NLin6Asym space. First, a reference volume and its skull-stripped version were generated using a custom methodology of fMRIPrep. Automatic removal of motion artifacts using independent component analysis (ICA-AROMA, Pruim et al. 2015) was performed on the preprocessed BOLD on MNI space time-series after removal of non-steady state volumes and spatial smoothing with an isotropic, Gaussian kernel of 6mm FWHM (full-width half-maximum). Corresponding “non-aggressively” denoised runs were produced after such smoothing. Additionally, the “aggressive” noise-regressors were collected and placed in the corresponding confounds file. All resamplings can be performed with a single interpolation step by composing all the pertinent transformations (i.e. head-motion transform matrices, susceptibility distortion correction when available, and co-registrations to anatomical and output spaces). Gridded (volumetric) resamplings were performed using antsApplyTransforms (ANTs), configured with Lanczos interpolation to minimise the smoothing effects of other kernels (Lanczos 1964). Non-gridded (surface) resamplings were performed using mri_vol2surf (FreeSurfer).

#### Functional data preprocessing

For each of the 8 BOLD runs found per subject (across all tasks and sessions), the following preprocessing was performed. First, a reference volume and its skull-stripped version were generated using a custom methodology of fMRIPrep. Head-motion parameters with respect to the BOLD reference (transformation matrices, and six corresponding rotation and translation parameters) are estimated before any spatiotemporal filtering using mcflirt (FSL 6.0.5.1:57b01774, Jenkinson et al. 2002). The estimated fieldmap was then aligned with rigid-registration to the target EPI (echo-planar imaging) reference run. The field coefficients were mapped on to the reference EPI using the transform. BOLD runs were slice-time corrected to 0.451s (0.5 of slice acquisition range 0s-0.902s) using 3dTshift from AFNI (Cox and Hyde 1997, RRID:SCR_005927). The BOLD reference was then co-registered to the T1w reference using mri_coreg (FreeSurfer) followed by flirt (FSL 6.0.5.1:57b01774, Jenkinson and Smith 2001) with the boundary-based registration (Greve and Fischl 2009) cost-function. Co-registration was configured with six degrees of freedom. Several confounding time-series were calculated based on the preprocessed BOLD: framewise displacement (FD), DVARS and three region-wise global signals. FD was computed using two formulations following Power (absolute sum of relative motions, Power et al. (2014)) and Jenkinson (relative root mean square displacement between affines, Jenkinson et al. (2002)). FD and DVARS are calculated for each functional run, both using their implementations in Nipype (following the definitions by Power et al. 2014). The three global signals are extracted within the CSF, the WM, and the whole-brain masks. Additionally, a set of physiological regressors were extracted to allow for component-based noise correction (CompCor, Behzadi et al. 2007). Principal components are estimated after high-pass filtering the preprocessed BOLD time-series (using a discrete cosine filter with 128s cut-off) for the two CompCor variants: temporal (tCompCor) and anatomical (aCompCor). tCompCor components are then calculated from the top 2% variable voxels within the brain mask. For aCompCor, three probabilistic masks (CSF, WM and combined CSF+WM) are generated in anatomical space. The implementation differs from that of Behzadi et al. in that instead of eroding the masks by 2 pixels on BOLD space, a mask of pixels that likely contain a volume fraction of GM is subtracted from the aCompCor masks. This mask is obtained by thresholding the corresponding partial volume map at 0.05, and it ensures components are not extracted from voxels containing a minimal fraction of GM. Finally, these masks are resampled into BOLD space and binarized by thresholding at 0.99 (as in the original implementation). Components are also calculated separately within the WM and CSF masks. For each CompCor decomposition, the k components with the largest singular values are retained, such that the retained components’ time series are sufficient to explain 50 percent of variance across the nuisance mask (CSF, WM, combined, or temporal). The remaining components are dropped from consideration. The head-motion estimates calculated in the correction step were also placed within the corresponding confounds file. The confound time series derived from head motion estimates and global signals were expanded with the inclusion of temporal derivatives and quadratic terms for each (Satterthwaite et al. 2013). Frames that exceeded a threshold of 0.5 mm FD or 1.5 standardized DVARS were annotated as motion outliers. Additional nuisance timeseries are calculated by means of principal components analysis of the signal found within a thin band (crown) of voxels around the edge of the brain, as proposed by (Patriat, Reynolds, and Birn 2017). The BOLD time-series were resampled into standard space, generating a preprocessed BOLD run in MNI152NLin6Asym space. First, a reference volume and its skull-stripped version were generated using a custom methodology of fMRIPrep. Automatic removal of motion artefacts using independent component analysis (ICA-AROMA, Pruim et al. 2015) was performed on the preprocessed BOLD on MNI space time-series after removal of non-steady state volumes and spatial smoothing with an isotropic, Gaussian kernel of 6mm FWHM (full-width half-maximum). Corresponding “non-aggressively” denoised runs were produced after such smoothing. Additionally, the “aggressive” noise-regressors were collected and placed in the corresponding confounds file. All resamplings can be performed with a single interpolation step by composing all the pertinent transformations (i.e. head-motion transform matrices, susceptibility distortion correction when available, and co-registrations to anatomical and output spaces). Gridded (volumetric) resamplings were performed using antsApplyTransforms (ANTs), configured with Lanczos interpolation to minimise the smoothing effects of other kernels (Lanczos 1964). Non-gridded (surface) resamplings were performed using mri_vol2surf (FreeSurfer).

Many internal operations of fMRIPrep use Nilearn 0.9.1 (Abraham et al. 2014, RRID:SCR_001362), mostly within the functional processing workflow. For more details of the pipeline, see the section corresponding to workflows in fMRIPrep’s documentation. Copyright Waiver

*The above boilerplate text was automatically generated by fMRIPrep with the express intention that users should copy and paste this text into their manuscripts unchanged. It is released under the CC0 licence*.

## Statistical Analyses

### Analysis of behavioural data

Performance was measured during the training tasks as the percentage of correct responses in the test phases (see above). On the first day, this corresponded to a total of 720 trials (change level of 50%). Performance on unseen numbers was measured using the two extended test phases after 20 and 40 blocks, which consisted of 48 trials. On the second day, participants were presented with 180 test trials on learned numbers and 24 test trials consisting of previously-unseen numbers.

In the recall task, performance was measured relative to probe trials, of which there were 16 per run (therefore 64 in the eyetracker and 128 in the fMRI scanner; chance level 50%).Participants had slightly longer to reply in the fMRI scanner (1.4s instead of 1.2s).

Half of the probe trials consisted of previously-unseen numbers. To compare performance on seen and unseen numbers, a two-sided paired t-test was used with an alpha level of 0.05.

Differences in reaction time in the recall task were measured using repeated measures ANOVA (alpha of 0.05) to ask if state identity predicted the mean reaction time for each state, for numbers followed by a probe trial. This was particularly relevant for the second day, to ensure differences in reaction times (and thus potentially task difficulty) could not explain our model of interest. Post-hoc pairwise comparisons were then carried out using paired t-tests across combinations of states, using a two-tailed test with a Bonferroni-corrected alpha of 0.0083 to reflect six pairwise comparisons.

Questionnaire data was read by the experimenter to ensure no participants were aware of the experimental question. To obtain a measure of strategy, the percentage (of 57) who replied ‘memory’, ‘calculation’ or ‘both’ in the multiple choice question “How did you resolve the task today?” was measured.

### Representational similarity analysis

Representational similarity analysis (Kriegeskorte et al., 2008) is a correlational method which compares the (dis)similarity of neural signals in different conditions or trials to a matrix of predicted model-based distances.

To run RSA, we first ran a first-level GLM on the preprocessed data in order to extract beta values for each condition. The GLM included motion regressors from fmriprep (see above), as well as an intercept. We also included regressors modelling visual information displayed on the screen and motor actions, specifically separate regressors for each type of visual presentation (stimulus presentation, probe trial presentation, response presentation, fixation cross) and for left/right button presses. As conditions, we included the state identity for each presented number, modelled as a stick function.

This model was run separately on each of the 8 functional runs in native (subject) space (T1w), to obtain a map of beta values for each condition, run and every subject. This allowed us to run cross-validated dissimilarity measures, which identify signal common to the same condition across runs and thereby reduce the effect of run-wise noise. The cross-validated Mahalanobis distance (’crossnobis’; Diedrichsen et al., 2016; Walther et al., 2016) was used, using the residuals from the first-level GLM to estimate the relevant noise covariance matrix for each run.

As the main hypothesis concerned a specific region (namely, the entorhinal cortex), a region-of-interest (ROI) approach was used. ROIs were sourced from the Jülich Atlas (Amunts et al., 2020, version 3.0.3) in MNI space, taking the maximum probability maps (MPMs) for each region (right entorhinal cortex, left entorhinal cortex). ROIs were then mapped into subject space using ANTs and the transformation matrices output by fmriprep. The resulting ROIs had the following sizes: left entorhinal cortex (mean 111.5 voxels, standard deviation 12.3), right entorhinal cortex (mean 140.1 voxels, standard deviation 22.0).

Based on this, we then calculated a neural dissimilarity matrix (see above) using the four different conditions in the experiment. This was then compared to model matrices (see below) using a tie-corrected spearman rank correlation, resulting in a single correlation value. Partial correlation (using recursion) was run to account for the effect of other models. In order to perform group-level inference, the resulting correlation values for each participant were Fisher z-scored (to approximate normality) and tested against a null hypothesis of mean 0 using a one-sample t-test. To account for multiple hemispheric comparisons, a Bonferroni-corrected alpha level of 0.025 was used to determine statistical significance in our main affordance model.

### Representational similarity analysis: Searchlight

Searchlight analysis was run following the procedure outlined in (Kriegeskorte et al., 2006). The logic of this analysis is that representational similarity analysis can be run within spheres to provide a whole-brain map of multivariate responses to different models.

As before, all analyses were carried out at subject level. Spheres were generated using rsatoolbox for Python v0.2.0 (Nili et al., 2014), with a radius of 3 voxels (7.5mm) and a minimum number of surrounding voxels necessary set as 30%. This analysis was run at a whole-brain level.

We followed the exact same approach as for the ROI-based RSA analysis; that is, neural dissimilarity matrices were calculated using crossnobis distance (with residuals from the first-level GLM for noise covariance calculation), and then compared to model matrices using a tie-corrected spearman correlation prior to z-scoring. We excluded the effect of other models using partial correlation.

In order to run group-level analyses, the resulting maps were then converted back into MNI space, and TFCE-based cluster correction (Smith & Nichols, 2009) was run within the whole brain. No significant clusters were found at a whole-brain level.

Our main model of interest concerned the similarity of states in terms of whether they allowed the same or different actions. To test this, we used a count-based dissimilarity matrix which indicated the number of shared actions between states (subtracted from the maximum, 2). For example, as states 1 and 4 both allow the same two actions (−2, +2) their dissimilarity is 0; states 1 and 2 only have one shared action (+2), so their predicted dissimilarity is 1. The count-based approach was also used for the two other control models, link distance (the minimum number of steps between two states), and action magnitude (the number of shared actions, independently of the numerical sign).

### Analysis of eyetracking data

The eyetracking data was first cleaned by removing blinks and surrounding data (using a window of 100 ms) from the time series. Data was then segmented into trials using a window of -500ms to 2500ms around stimulus onset. Trials with less than 700ms of data remaining were excluded from analyses. Trials of each condition were then grouped together and the mean fixation in x and y was taken for each timepoint. To account for temporal noise, the data was then smoothed with a Gaussian kernel with a standard deviation of 4.25ms (roughly corresponding to a full width half maximum kernel width of 10ms) and a maximum window of 20ms within each condition and subject. This resulted in a single time series in both x and y for each condition, for each subject.

To test for an action-based change in gaze behaviour, we compared the directional shift as a function of different states. State 2 allows only positive operations, while state 3 allows only negative operations. We obtained the relative shift by subtracting fixations for state 3 from state 2 in both the x and y axis, for every time point. If the relative position difference is positive, this means an eye movement further right or up for states which afford positive actions relative to states which afford negative actions.

In order to carry out group-level analyses, we ran a temporal cluster correction using the MNE package (Gramfort et al., 2013) based on a 1-sample t-test with 10,000 permutations. As the hypothesis predicted a positive movement, a one-tailed test was used with an alpha level of 0.025 to account for multiple comparisons across the two axes (Bonferroni correction).

Finally, we compared the results to performance. To do so, for each participant we identified the median change in gaze position across every timepoint within all previously-identified significant clusters. This was carried out both in x and y, to give a single value in each axis which reflected how strongly the directional effect was for a given participant within the cluster. For both analyses, a more positive value indicated a stronger effect. This was then correlated with performance in probe trials (pearson r) in the first day to obtain a measure of how much gaze effects reflected individual performance. An alpha level of 0.025 was used to account for tests on each axis (Bonferroni).

## Acknowledgements

S.T.’s research is supported by a Minerva Fast Track Fellowship of the Max Planck Society. CD’s research is supported by the Max Planck Society and the Kavli Foundation. A.E. is enrolled in the International Max Planck Research School on Neuroscience of Communication and a doctoral programme in Cognitive and Brain Sciences at the University of Trento. This work was supported by the European Research Council (ERC-StG, NOAM 804422) and the Italian Ministry of University and Research (MUR-FARE, MODGET R18WJMSNZF) attributed to R.B. We thank all scanner technicians and research assistants for their support in participant recruitment and data collection, as well as the other members of our labs for feedback and code sharing, in particular Simone Viganò, Yangwen Xu and Rebekka Tenderra for advice regarding fMRI design and analyses.

## SUPPLEMENTARY MATERIALS

### Test of noise in cross-validated RSA

As crossnobis is a cross-validated distance measure, we can also use the resulting similarity scores to assess how much a given region is consistently responsive to the conditions across runs. Namely, a score greater than 0 indicates a consistent response (Diedrichsen et al., 2016; Walther et al., 2016). To test this, for each region we calculated the mean distance within the neural RDM for each ROI and tested against 0 using a Wilcoxon signed rank test. Note that this is a conservative test, as we predict a distance of 0 for some state combinations in the affordance model.

Both regions showed mean distance values greater than 0 (Wilcoxon signed rank test, p<0.0001 for both ROIs), indicating the results were driven by signal (that is, consistent voxel patterns across runs) rather than noise.

It is worth noting, furthermore, that if only subjects with the mean distances greater than 0 are selected, the affordance RSA effect in the right entorhinal cortex becomes stronger (n=35, t(34)=3.15; p=0.0046 in a one-tailed t-test), whereas in the left entorhinal cortex this is not the case (n=40, t(39)=-0.0090; p=0.50). This further indicates that for subjects with consistent neural pattern responses between runs for each condition (i.e. BOLD responses explained by conditions of our GLM, not condition-irrelevant noise), the right entorhinal cortex specifically represents conceptual affordances.

**Supplementary Figure 1:**
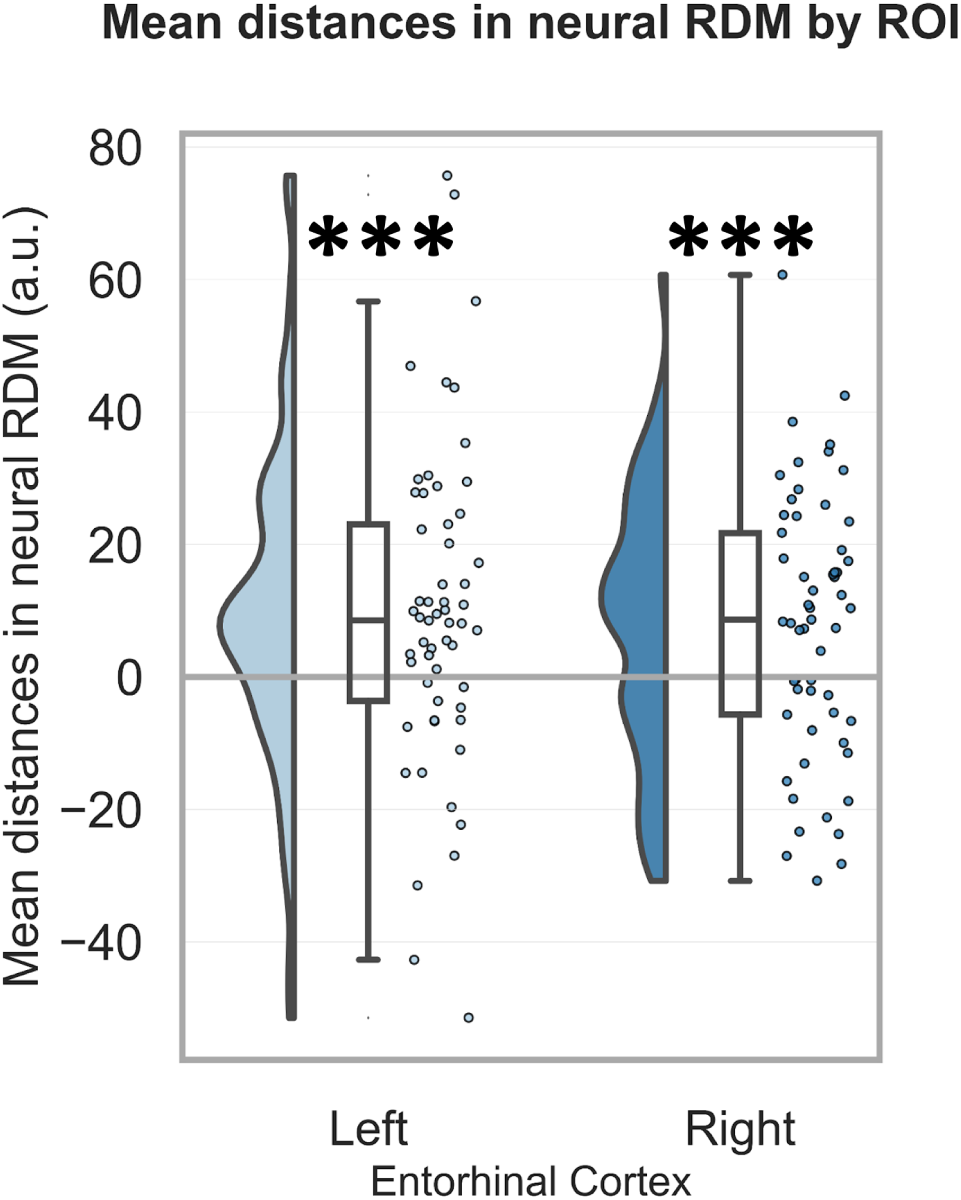
Cross-validated distance measures produce distance values greater than 0 if there are similar voxel patterns across runs. This allows testing for noise within specific ROIs. In both ROI used, we find mean distance values across all cells of the RDM matrix were greater than 0 (Wilcoxon signed-rank test, p<0.0001).

### Reliability-based voxel selection

To ensure that the main effect was not driven by noise at the voxel level, we implemented a modified version of reliability-based voxel selection (Tarhan & Konkle, 2020). The initial method was designed for condition-rich designs, and the authors advise caution in multivariate analyses due to potential imbalances. As the trials were balanced across conditions, we considered that the method was appropriate for RSA. Nonetheless, as the design had only four conditions, it was necessary to adapt the method to be sure that it was robust.

To do this, we considered a permutation-based approach. Rather than simply correlating betas in averaged halves of the data across conditions (as with the original method), which would involve a 4-by-4 correlation, we chose to (1) use Euclidean distance between vectors of beta values, which is more robust than correlation for small vectors, and (2) shuffle the data across conditions for a given voxel (1000 times), and re-calculate resultant distances to obtain an estimate of how likely a given Euclidean distance was based on the distribution of possible distances. Using this approach, we could assign a probability value to each voxel which described how much of the distribution was less than the actual, non-shuffled distance measure. A low probability value indicates a consistent response for a given voxel for each condition across runs.

The resultant probability maps were intersected with entorhinal ROIs to generate smaller ROIs within which voxels were more likely to be condition-responsive. Checks on simulated data confirmed that this procedure did not increase the rate of false positives. Nonetheless, this analysis assumes that the brain does indeed represent conditions differently, so it should be noted that results using this method should be interpreted as “[non] significant given that the brain region of interest is differentially responsive to the experimental conditions”. As such, we consider these analyses confirmatory rather than a primary test of the main hypotheses.

As the probability values were typically quite high, we chose a liberal threshold of p<0.2 for the intersected ROIs (resulting in small ROIs: right entorhinal cortex mean voxels 31.2, std. 17.4; left entorhinal cortex mean voxels 22.3, std. 10.4). Despite the much smaller regions-of-interest, we still found a significant affordance effect in the right entorhinal cortex (t(56)=2.30; p=0.0126), suggesting the main finding is not the result of variations in voxel-level noise captured within anatomical regions-of-interest (left entorhinal cortex: affordance: t(56)=0.75; p=0.227).

**Supplementary Figure 2:**
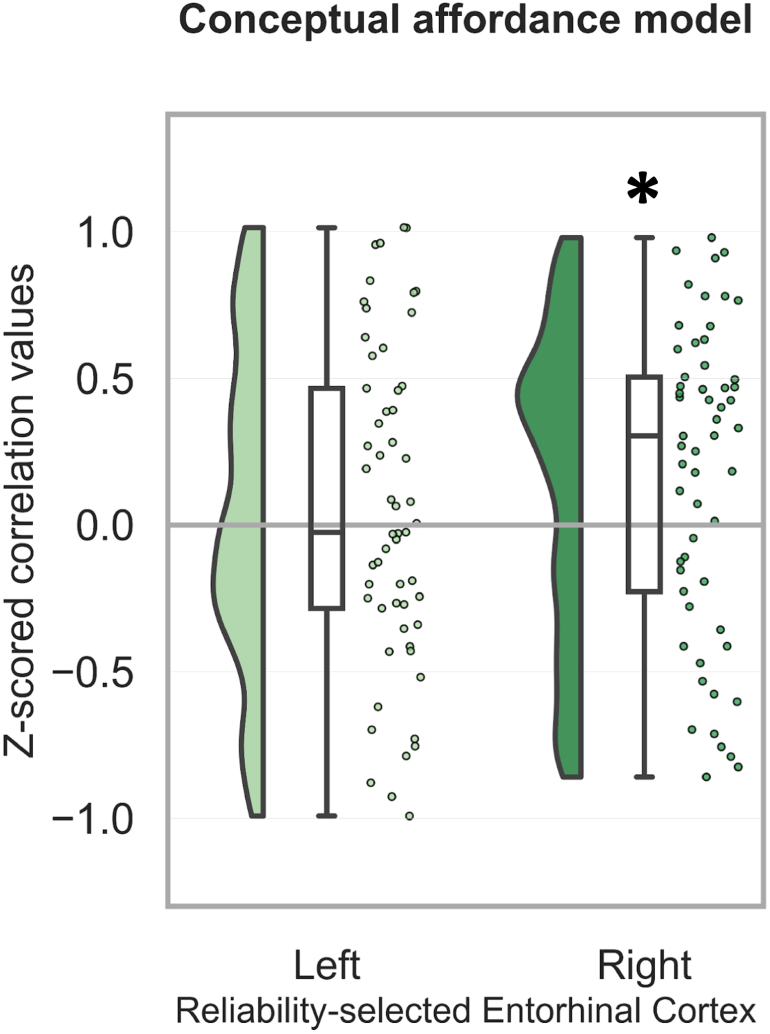
Using a variant of reliability-based voxel selection, we created smaller regions-of-interest in which neural responses were more consistent, for each voxel, across all eight runs. Within these reduced ROIs, the main effects found in the primary analysis were still present, further confirming that our results are driven by signal rather than noise.

### Comparison of affordance effect in right entorhinal cortex to noise ceiling

To quantify the possible correlation of an ideal model with the right entorhinal cortical signal, we used inbuilt functions from rsatoolbox to quantify the noise ceiling (Nili et al., 2014; Schütt et al., 2023). The affordance model was not significantly below the noise ceiling (p=0.246). This indicates that the affordance effect was as strong as possible given experimental noise.

**Supplementary Figure 3:**
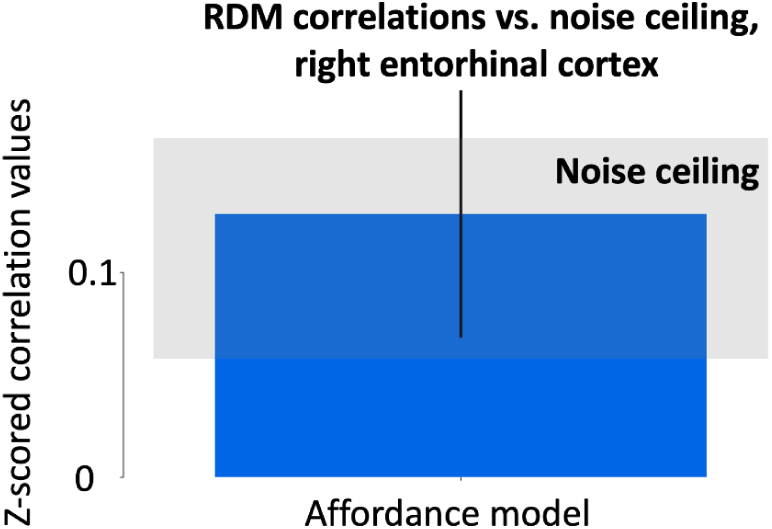
The affordance model effect was within the noise ceiling (grey bar), indicating that the effect was as strong as possible given experimental noise and condition-wise responses.

